# TOCA-2 regulates gonad development in *C. elegans*

**DOI:** 10.64898/2026.04.26.720826

**Authors:** Yogesh Pratap, Tanushree Sinha, Anup Padmanabhan

**Affiliations:** Department of Biology, Ashoka University, No-2 Rajiv Gandhi Education City, Sonipat, Haryana India, 131029

## Abstract

The development of *C. elegans* gonad involves the coordinated action of diverse biochemical factors and physical forces. The precise roles and interconnections of these diverse components remain poorly understood. TOCA-2, the *C. elegans* ortholog of mammalian TOCA-1 (Transducer of CDC42 Dependent Actin Assembly) is an F-BAR domain protein known to play important roles in oocyte maturation and embryogenesis through membrane remodelling and regulation of actin cytoskeleton. Here we demonstrate that TOCA-2 is actively involved in maintaining the structural integrity and function of the *C. elegans* gonad. *toca-2(null)* animals exhibit pronounced architectural defects with particularly strong perturbations on the dorsal side of the gonad. Phalloidin staining and cytoplasmic particle velocity analysis revealed that the actomyosin corset surrounding the common rachis in the syncytial germline is severely disorganized in the mutants. This disorganization leads to significant disruptions in cytoplasmic flow across different regions of the syncytium. Together, our findings quantitatively highlight mechanisms underlying gonad morphogenesis and maintenance, establishing TOCA-2 as a key regulator of these processes. This work also provides a framework for positioning TOCA-2 within broader biochemical pathways governing organogenesis and other developmental processes dependent on actomyosin dynamics in *C. elegans*.

## Introduction

The *C. elegans* gonad has been extensively used as a model to study organogenesis from multiple perspectives. This includes, identifying the role of collective cell migration in tissue morphogenesis [1]; exploring the interaction between cells and the decision-making process within cells to unravel the mechanical regulation of specialized structures [1][2]; and understanding the role of actomyosin contractility at various scales in regulating tissue structural organization [3]. The *C. elegans* hermaphrodite gonad comprises of two symmetrically arranged U-shaped tubes [5], one arm of the gonad is arranged above the intestine and another is below the intestine providing a ‘hugging morphology’ [6]. Although the intestine in *C. elegans* develops during early embryogenesis, the gonad begins to develop post embryonically [7][1]. Distal end of both the arms has somatic DTC (Distal Tip Cells) which release metalloproteases and helps in gonad elongation [8]. At L1 and L2 stages the gonad arms elongate opposite to each other along the ventral side. During the early L3 stage, both the arms take a U-turn at different Z-plane to maintain the hugging morphology with the intestine. These turns are carried out through a torque generated because of asymmetric DTC-matrix adhesion[8][3][1]. At L4, gonadal arms start approaching each other along the dorsal side. The young adult stage marks the onset of reproductive function [1]. The dorsal sides of the gonad house germ cells within a syncytium. These germ cells are partially open towards the common cytoplasm known as rachis [9][3]. Cytoplasmic flow from the dorsal to the ventral side, beginning during the young adult stage enables oocyte packaging and maturation. These oocytes then enter the spermatheca, and following fertilization, they exit into to the uterus [10][11]. A contractile network resembling a corset at the tissue level surrounding the rachis, composed of actomyosin, serves a crucial function in preserving the integrity of the syncytial germline structure. This Is achieved by exerting forces in two perpendicular directions, facilitating cytoplasmic flow, regulating membrane tension, and inwardly pulling the germ cell membranes radially (Fig. 1a) [3].

**Figure 1.**
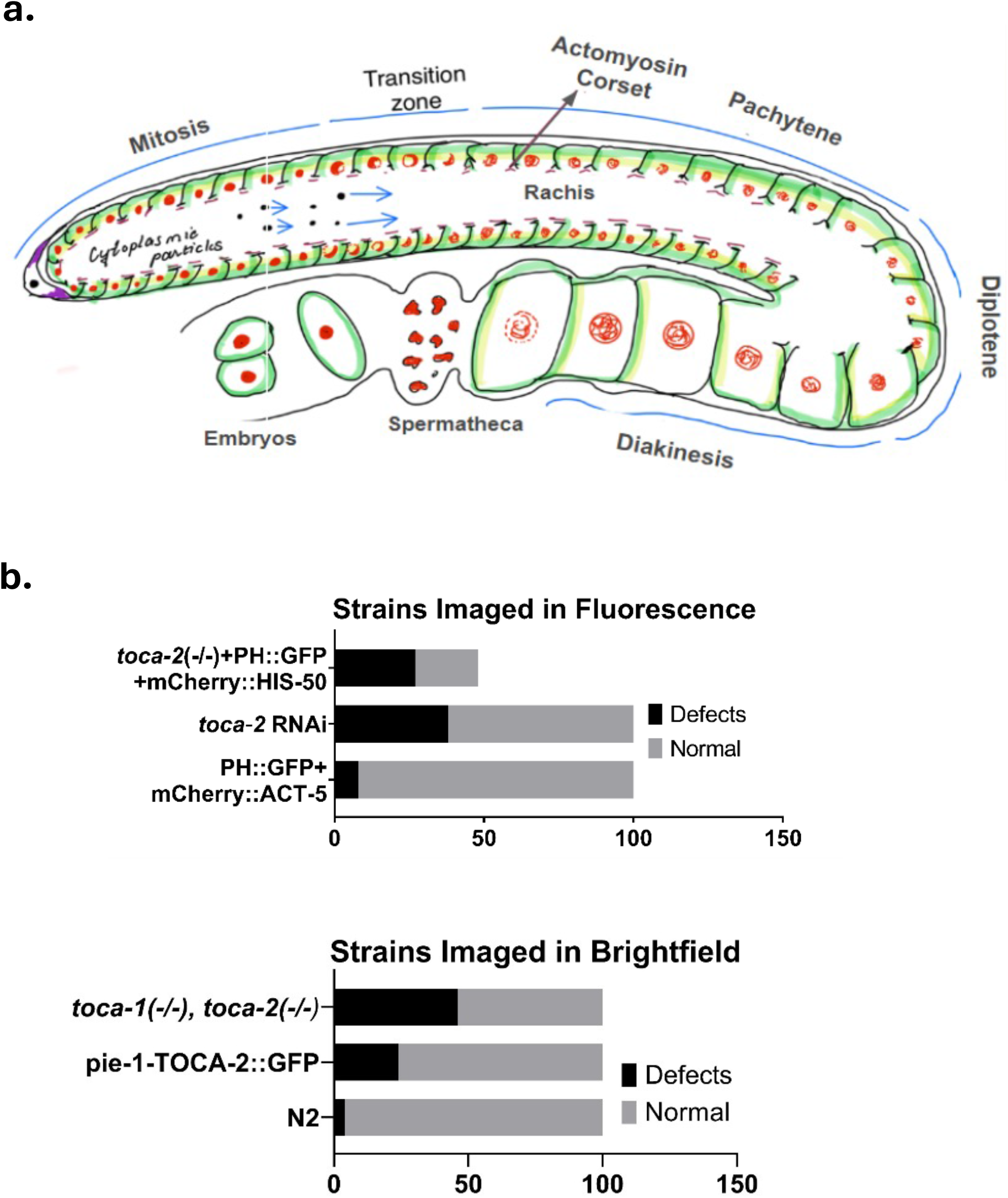
a. Schematic representation one of the two arms constituting the gonad of an adult hermaphrodite *C. elegans*. b between the worms with gonad structural defect and worms having normal morphology in TOCA-2 mutants. The total worms screened were 100 in all the mutants except toca-2 (-/-) + PH::GFP + mCherry::his-58 which only had 48 worms.

Development of the *C. elegans* hermaphrodite gonad involves collective cell migration and coordinated differentiation, which extensively requires regulation of the cytoskeleton and dynamic reorganization of the plasma membrane. The TOCA family of proteins has been implicated in regulating cytoskeleton-membrane interactions through their specialized domains. In *C. elegans*, the non-essential TOCA-2 (hereafter TOCA-2) contains a curvature-sensing F-BAR domain at the Nterminal, an HR1 domain that binds to CDC-42, and an N-WASP binding SH3 domain at the C-terminal [12](Fig. S1). Previous reports demonstrated TOCA-2 localizes to rachis membranes, indicating a possible actin-dependent role in gonad morphogenesis and oocyte development [12][14]. Giuliani et. al. demonstrated that disruption of clathrin-mediated endocytosis in TOCA-2 mutants hinders intestinal yolk uptake by the germline, affecting oocyte maturation and reduced brood size. Additionally *toca-2(null)* mutants also show statistically significant embryonic lethality due to the *gex* (*g*ut on the *ex*terior) defect [12][13].

In this study, we investigated the role of TOCA-2 in gonad morphogenesis in *C. elegans*. We show TOCA-2 depletion results in defective in actin organization in thel gonad syncytium, disrupting cytoplasmic flow leading to deformed gonad architecture. Tissue specific expression of TOCA-2 in the germline is sufficient to rescue germline deformation. In addition to previously reported roles of TOCA-1/2 in endocytosis of yolk granules, these findings highlights the importance of TOCA-2 in organogenesis and tissue mechano-regulation of *C. elegans* gonad.

## MATERIALS AND METHODS

### Growth and maintenance of strains

*C. elegans* and bacterial strains used in this study are listed in Table S1. Primers used to confirm deletion mutants in *toca-1* and *toca-2* are listed in Table S2. All *C. elegans* strains were maintained at 20° C on Nematode Growth Medium (NGM) agar plates seeded with *E. coli* OP50 (Stiernagle, 2006). All bacterial cultures were grown in Luria-Bertani (LB) broth at 37°C at 180 rpm.

### RNA interference

RNA interference was carried using feeding of the *E. coli* strain HT115 (DE3) expressing L4440 plasmid into which sequences targeting *toca-2* was cloned. HT115 (DE3) containing RNAi clones was cultured in LB broth containing (100 μg/mL) and tetracycline (12.5 μg/mL) at 37° C and seeded on to the NGM plate containing 1mM isopropyl β-D-thiogalactoside (IPTG) and 100 μg/mL ampicillin as described previously(Kamath, 2003). L4440 (vector alone) was used as the RNAi control. F2 embryos from animals grown on *toca-2(RNAi)* plates were isolated and allowed to hatch on plates devoid of bacteria. L1-stage synchronized worms were subsequently transferred to *toca-2(RNAi)* plates for analyzing post embryonic development of germline architecture.

### Microscopy

The screening of the young adult worm’s gonad architecture was conducted using an Olympus BX63 Upright epi-fluorescence microscope. A total of 100 worms were examined from both the mutant and control groups (N2). The worms were mounted on a 3% agarose pad to prevent desiccation, and 0.05% levamisole was used to anesthetize them. Image acquisition was controlled by Olympus CellSens Dimension software. Gonads that exhibited significant structural deviations from the normative descriptions were classified as defective. Comparison of the ratio of the number of worms with defective and normal gonad architecture was performed in GraphPad Prism 9.0. The Mann-Whitney test and Student’s t-test was employed in case of non-normal and normal distributed data, respectively.

### Mitotic Zone Analysis

Worms washed in M9 buffer and anesthetized in 0.025% levamisole were dissected near the pharynx using a Dispovan 31G, 1ml syringe. Extruded gonads were fixed in 2% PFA (Paraformaldehyde) for 15 minutes. The sample was imaged under an Olympus IX83 Inverted Microscope (Spinning-disc confocal) and excited at 488 and 561 nm laser lines using an OBIS Coherent laser system. The length of the mitotic zone on the distal side of the gonad was measured from the distal tip cell (DTC) to the transition zone, characterized by two or more crescent-shaped nuclei in a row. Each measurement had three replicates.

### Particle Image Velocimetry (PIV)

PIV for cytoplasmic flow in distal arm of the gonad was carried out through ImageJ plugin. The time-lapse DIC video was recorded for 4 minutes with a 2-second gap between consecutive frames. Custom macros and Python codes were used to automate image processing and analysis (codes in appendix). First-pass PIV with an interrogation window size of 32×32 pixels and 50% overlap was employed on an ROI of size 30.3×28.3 microns. The time-averaged velocities were calculated by averaging the instantaneous velocities of one interrogation window over 120 pairs of consecutive frames, and the spatial velocity was calculated by averaging instantaneous velocities of all interrogation windows over one pair of consecutive frames.

## Results

### TOCA-2 depletion leads to defects in gonad architecture

To determine whether somatic depletion of TOCA-2 affects the organization of the *C. elegans* germline, we expressed TOCA-2 fused to GFP at the C-terminus under the germline -pecific *pie-1* promoter (*pie-1*::TOCA-2::GFP) in a *toca-2(null)* background. Microscopic analysis revealed that 24% of mutant animals exhibited structural and positional defects in the gonad, compared to only 4% in the control animals indicating significant germline abnormalities upon TOCA-2 depletion. Given that the gonad occupies a significant fraction of the worm’s body volume, changes in gonad morphology are expected to influence overall body shape. Accordingly, we performed a time-course analysis of body width. The thickness (w) of control and TOCA-2-depleted animals expressing TOCA-2::GFP in germline, followed the quadratic relationship: w(control, µm) = 0.01t^2^ − 0.46t + 19.57 and w(*toca-2(null)*^pie-1::TOCA-2^, µm) = 0.01t^2^ − 0.26t + 16.14, respectively (Fig. S2). To further characterize these defects, we performed live fluorescence imaging using strains co-expressing PH::GFP (membrane marker) and ACT-5::mCherry (localized to the apical side of the intestine). A quantitative analysis of worms depletion of TOCA-2 revraled a ∼30% increase in animals displaying gonadal defect (Fig. 1b). Efficient depletion was confirmed by loss of TOCA-2::GFP fluorescence (Fig. S3). Subsequent screening revealed a 30% increment in worms exhibiting gonad defects (Fig 1b).

*C. elegans* expresses two TOCA-2 paralogs – TOCA-1 and TOCA-2. To check for functional redundancy, we analysed gonad architecture in *toca-2(null); toca-1(null)* double mutants. Depletion of both TOCA-1 and TOCA-2 resulted in 42% of worms with gonad structural and positional defects compared to the wild type. Additionally, not only did the fraction of worms with defective gonads increase, but the defects also became more pronounced. To evaluate individual contributions from TOCA-1 and TOCA-2 we analysed *toca-1(null)* and *toca-2(null)* animals expressing PH::GFP. We notice that 27/48 *toca-2(null)* -worms exhibited significant structural defects in the gonad. This argues against possible synergistic effect between TOCA-1 and TOCA-2. Together, these results indicate that TOCA-2 plays a dominant role in maintaining gonad architecture in *C. elegans*.

### TOCA-2 mutants exhibit four classes of gonadal defects

We detected four predominant defects during our phenotypic analysis: (1) failure of the gonad to properly navigate around the intestine, leading to compression within a restricted region; (2) loss of directionality in the dorsal arm, resulting in aberrant intersections along the dorso-ventral (DV) axis; (3) distortion of the dorsal-to-ventral U-turn; and, (4) complete disassociation between the gonad and intestine, disrupting the characteristic “hugging’ morphology. Among these, the third defect was most frequent upon TOCA-2 depletion (Fig. S4). Quantification of the first two defect classes was performed by normalizing the gonad measurement to body width. Specifically, gonad compression was assessed by measuring the width of the affected region, while directional defects were quantified by measuring the distance between the dorsal arm and the body wall (Fig. 2a and 2e). The percentage error in gonad width measurement was determined to be 1.28µm, based on z-plane variability (Fig. S5). Overall, these analyses demonstrate that loss of TOCA-2 results in statistically significant alterations in gonad morphology.

**Figure 2:**
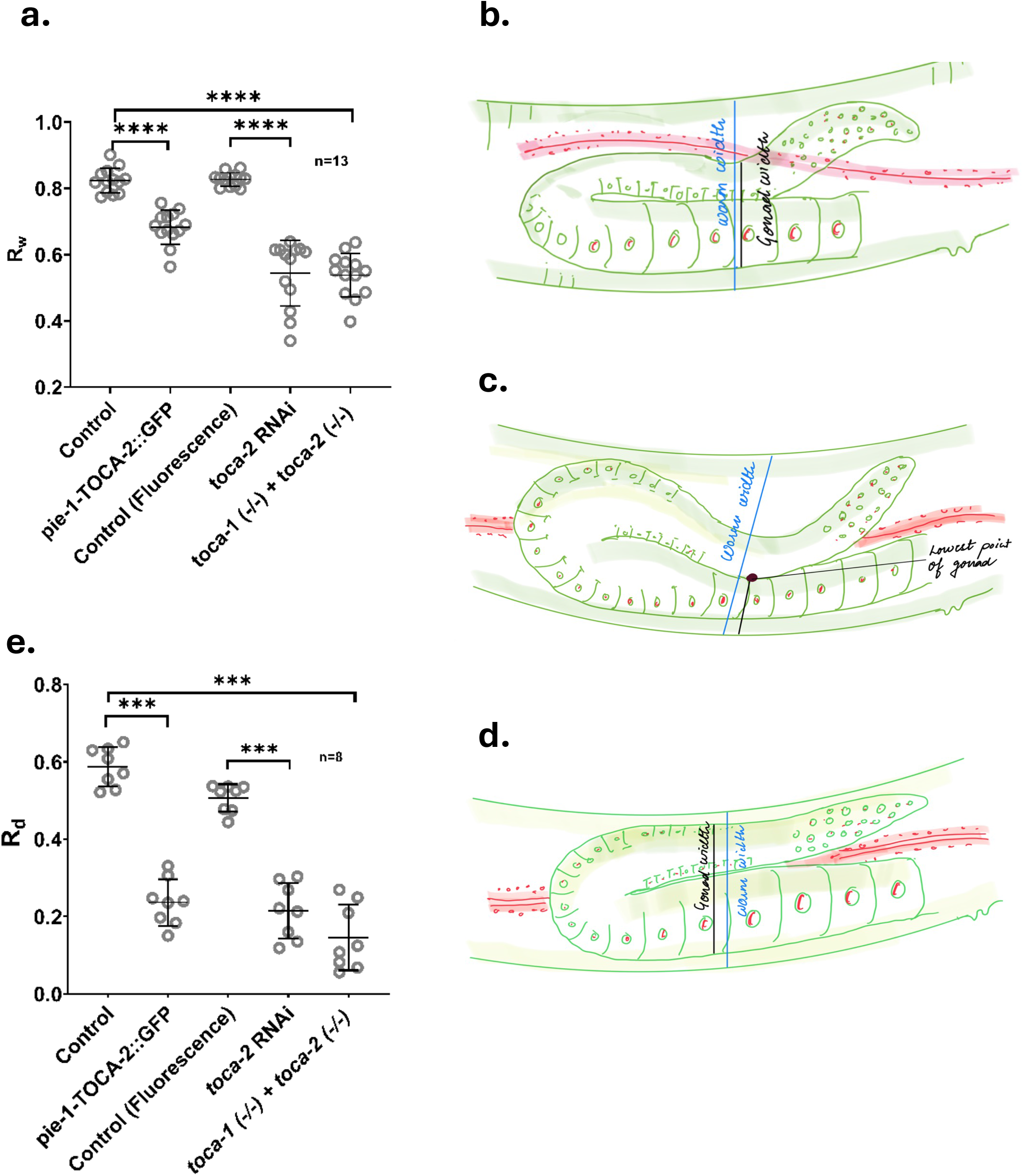
**a**. Comparing constricted gonad width in TOCA-2 mutants. Statistical analysis was conducted using the Mann-Whitney test, n=13. **b**,**c**,**d** Schematic diagrams illustrating the locations from which measurements were taken to quantify the defective gonads in the first and second defect categories, as well as in wild-type worms, respectively. e. Assessment of the crossing DV axis defect in TOCA-2 mutants, n=8

### Defects in gonad architecture arises ∼72 hours post-embryogenesis

Gonad morphogenesis is governed by multiple mechanical forces. These include forces exerted on the gonad during its elongation due to proliferative pressure of the germ cells, and an opposing “friction” generated due to the basement membrane around the gonad [1]. Additionally the torque responsible for turning of the gonad from ventral to dorsal side generated due to asymmetric DTC-ECM adhesion. Internal cytoplasmic flow (F) also exerts radial forces on the syncytium due to cytoplasmic flow. To investigate the temporal onset of defects, we tracked gonad development in control, and *toca-2(-/-)* animals co-expressing PH::GFP and mCherry::his-58. Gonads imaged at 24-hour intervals starting from the L1 stage. While gonad elongation demonstrated non-linear growth dynamics, animals depleted of TOCA-2 exhibited slower growth relative to controls. Interestingly,, between 72-96 hours post embryogenesis, mutant gonads showed a transient increase in length despite evident structural defects, eliminating differences with controls (Fig 3a, 3f, 3h). Between 96-120 hours, mutant gonads exhibited pronounced structural defects accompanied by a significant reduction in length (shrinkage) compared to control (Fig 3a, 3g, 3h). This phenotype resembles gonad atrophy described by de la Guardia et al., although in mutants it occurs much earlier (day 2 post-L4 in *toca-2(null)* vs. day 8 in control) [15]. Notably, shrinkage was predominantly restricted to the dorsal arm of the gonad (Fig. S6). Two possible explanation may account for these observations. First, transient increased proliferation between 72-96 hours could lead to overcrowding and mechanical crumpling. Alternately, loss of TOCA-2 may disrupt the actomyosin corset, altering cytoplasmic flow and thereby compromising structural integrity.

**Figure 3:**
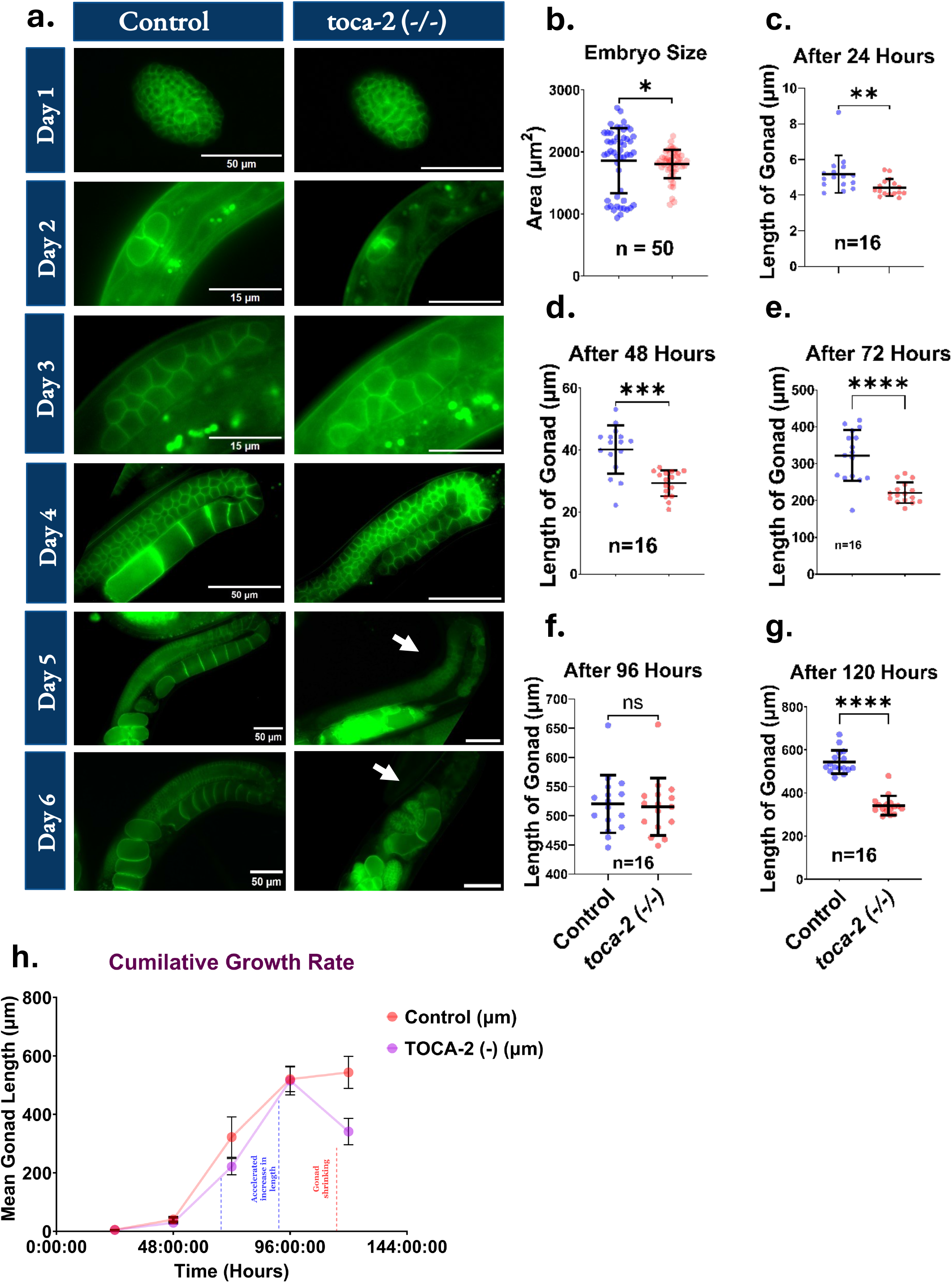
**a**. Micrographs depicting gonad development over time in both control and *toca-2(null)* mutants. **b**. Quantification of embryo size in control and *toca-2(null)* worms. Analyzed using the Mann-Whitney test. **c-g** Quantification of gonad length at various post embryonic developmental stages of control and *toca-2(null)* worms at 24-hour intervals (n=16). **h**. Mean length of the gonads plotted against time, with error bars representing standard deviation.

### TOCA-2 depletion does not alter mitotic zone length

To test whether increased proliferation contributes to gonad elongation, we measured the mitotic zone length in control and TOCA-2-depleted animals. In *C. elegans*, germ cells divide mitotically in the distal region before transitioning into meiosis proximally (Fig. 1a). We found no significant different in mitotic zone length between control and mutant animals (Fig. 4), suggesting that altered proliferation is unlikely to explain the transient increase in gonad length.

**Figure 4:**
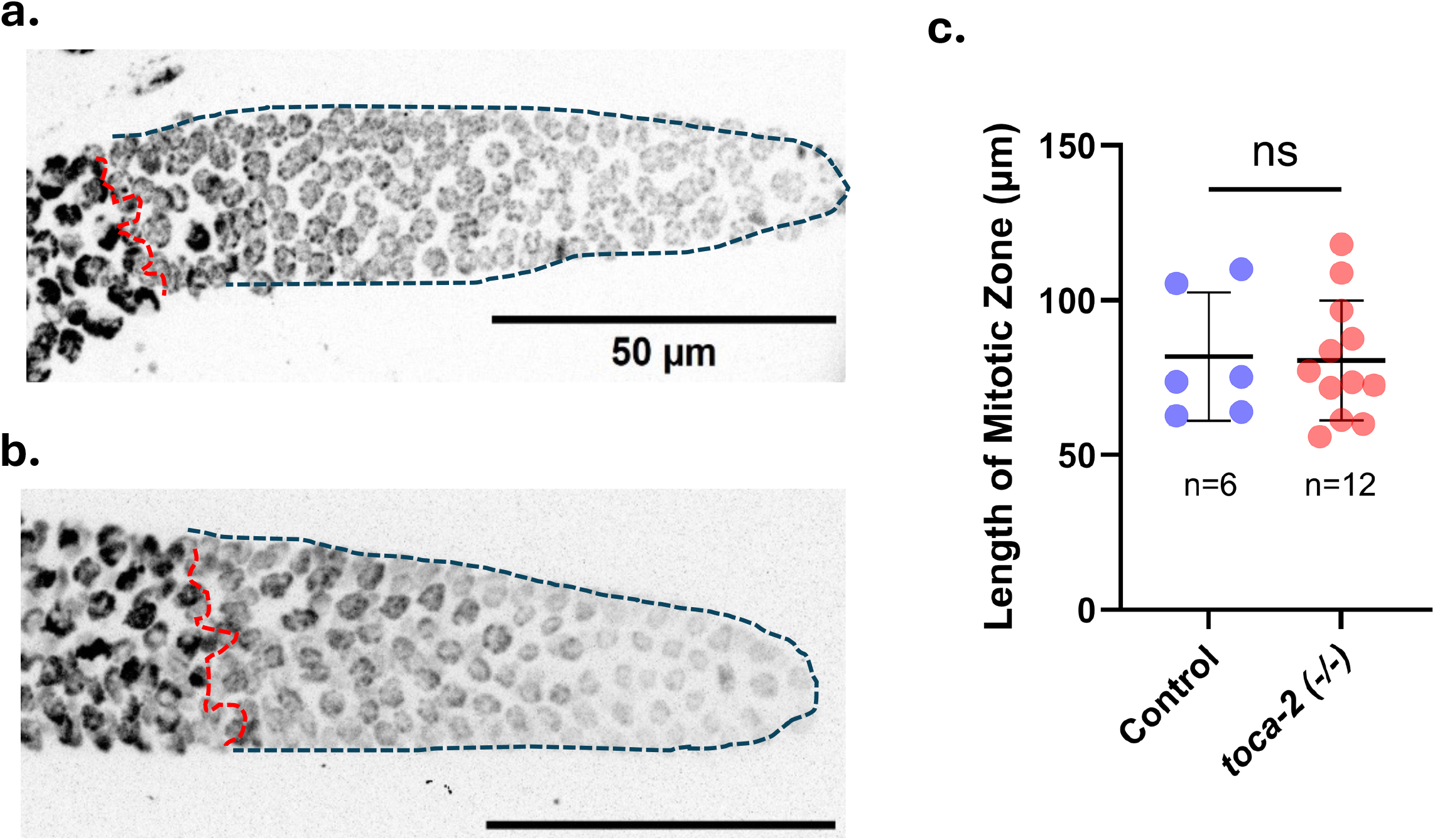
**a**,**b**. Micrographs representing mitotic zone in *toca-2(null)* and control gonads **c**. Quantification of mitotic zone in control (n= 6) and *toca-2(null)* (n= 12) worms. Mann-Whitney test for statistical comparison.

### TOCA-2 depletion disrupts the syncytium structure and results in disorganised cytoplasmic flow

TOCA-2 localizes to the partially ingressed membranes at the rachis and proteins with similar localization are known to regulate actin dynamics and syncytial organization [12][13]. Given the dorsal shrinkage phenotype, we examined actin organization using whole-worm phalloidin staining. Spinning disk confocal microscopy revealed disruption of the actomyosin corset in TOCA-2-depleted gonads. Unlike the uniform structure observed in controls, mutant syncytia exhibited irregular morphology with reduced width (Fig. 5a). Because cytoplasmic flows within is critical for oocyte development, we next performed Particle Image Velocimetry (PIV) analysis [12][13]. While mean instantaneous velocities were comparable between groups, velocity variability was significantly higher in *toca-2(null)* mutants ( standard deviation ∼8 µm/s vs. ∼5 µm/sin controls) (Fig. 5c and 5d). This likely reflects irregular rachis geometry, including dents and protrusions leading to fluctuating flow patterns. Time-average velocity maps further revealed altered flow distribution: higher velocities were observed near boundaries in *toca-2(null)* mutants, whereas in controls, velocities decreased smoothly from center to periphery (Fig 5e). Together, these findings demonstrate that loss of TOCA-2 disrupts actomyosin organization within the syncytium, leading to altered cytoplasmic flow and defective gonad architecture.

**Figure 5:**
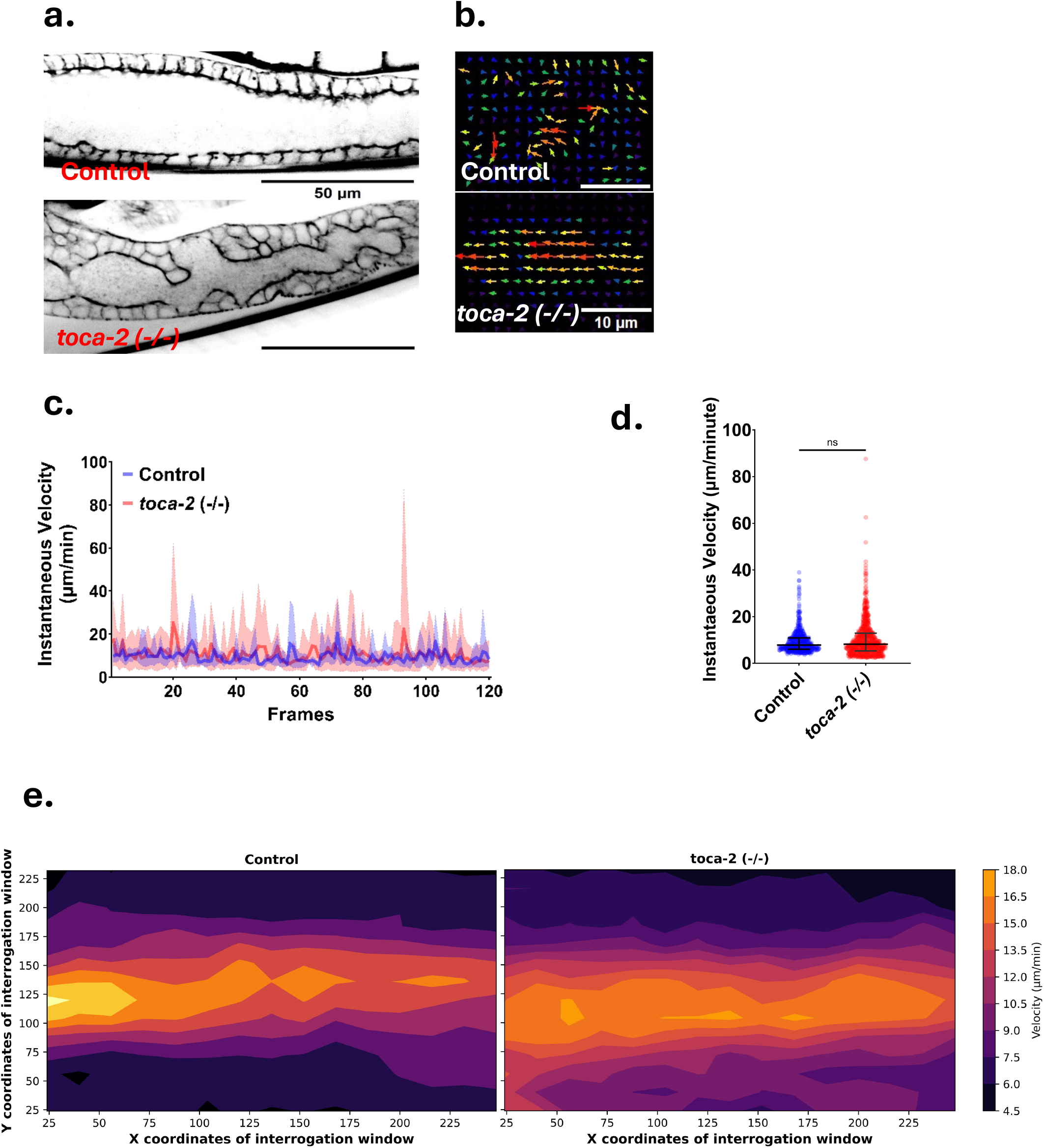
**a**. Confocal images of syncytium architecture in gonads from control and *toca-2(null)* mutants. b PIV analysis of the cytoplasmic flow in control and *toca-2(null)* mutant **c**,**e** Instantaneous velocities of all interrogation windows in consecutive frames *toca-2(null)* mutant (n=720) control (n=480). **d, e**, represents the contour plot of time average of instantaneous velocities across all the interrogation windows.

## Discussion

Several genes have been reported to regulate gonad architecture in *C. elegans*, including *ani-2, cdc-42* [1], *fli-1*, and *let-60* [12]; however, the role of the TOCA family proteins has remained unclear. In this study, we investigated how TOCA-2 cntributes to germline architecture. Our initial analysis revealed that 56% worms with *toca-2 (null)* exhibit significant gonadal defects. TOCA-2 is known to interact with CDC-42 through its HR1 domain[12]. Previous work by Watson et al. proposed that the mammalian ortholog of C. elegans TOCA-2, mTOCA-1, acts as a facilitator in the association between CDC-42 and N-WASP, leading to the recruitment and activation of ARP-2/3 [26]. Depletion of mTOCA-1, may reduce – but not abolish – the likelihood of CDC-42-N-WASP interaction. This provides a plausible explanation for why a substantial fraction (44%) of mutants still exhibit normal gonad morphology. Our spatiotemporal analysis indicates that gonadal defects begin to emerge ∼72 hours post-embryogenesis. By 96 hours, mutant gonads display pronounced crumpling accompanied by a reduction in length. We ruled our increased proliferation as the underlying cause, as mitotic zone length was comparable between control and *toca-2(null)* animals. Instead, whole-worm phalloidin staining revealed clear disruption of the actomyosin corset within the syncytium of *toca-2(null)* mutants. This structural disorganization correlates with altered cytoplasmic flow and likely contributes to defects in germline architecture.

The transient increase in gonad length between 72-96 hours remains an open question. Given that mitotic activity is unchanged, it is possible that mechanical forces contribute to this phenotype. Supporting this hypothesis, the rachis in *toca-2(null)* mutants appears narrower than controls (Fig. 5a), suggesting altered mechanical constraints or tension within the syncytium. Interestingly, actomyosin-dependent mechanical regulation is also central to spermathecal function in *C. elegans*, where repeated cycles of stretching and contraction facilitate oocyte fertilization and embryo extrusion. This process involves SPV-1, a RHO GTP-ase regulating F-BAR domain containing protein [18-20]. Given that TOCA-2 also contains an F-BAR domain, it is plausible that it similarly links membrane curvature sensing with actomyosin organization. Consistent with this in *toca-2(null)* mutants exhibit ∼11% embryonic lethality, altered spermathecal morphology (Fig. S7) and teratoma-like structures (Fig. S8)[21].

Given the conservation between *C. elegans* TOCA-2 and its mammalian orthologs, our findings may have broader implications for understanding organogenesis and cytoskeletal regulation in higher organisms. Dysregulation of actin dynamics is implicated in numerous disease, including cancer. Therefore, elucidating TOCA-2-mediated pathways could provide valuable insights into conserved mechanisms governing cytoskeletal organization and tissue architecture in development and disease.

## Acknowledgements

This work was supported by DBT-Wellcome India Alliance Fellowship (IA/I/18/1/503624) and ANRF Core research grant (CRG/2023/004638) to A.P, Department of Biotechnology Junior research Fellowship (DBT/2024-25/AshokaUni/2486) to T.S., and core funding support from the Trivedi School of Biosciences, Ashoka University. We acknowledge the infrastructure support from the Central Bio-imaging facility and Ashoka-Zeiss Core Imaging Facility at Ashoka University. Some strains were provided by the CGC, which is funded by NIH Office of Research Infrastructure Programs (P40OD010440).

## Supplementary Data

**Table S1.**
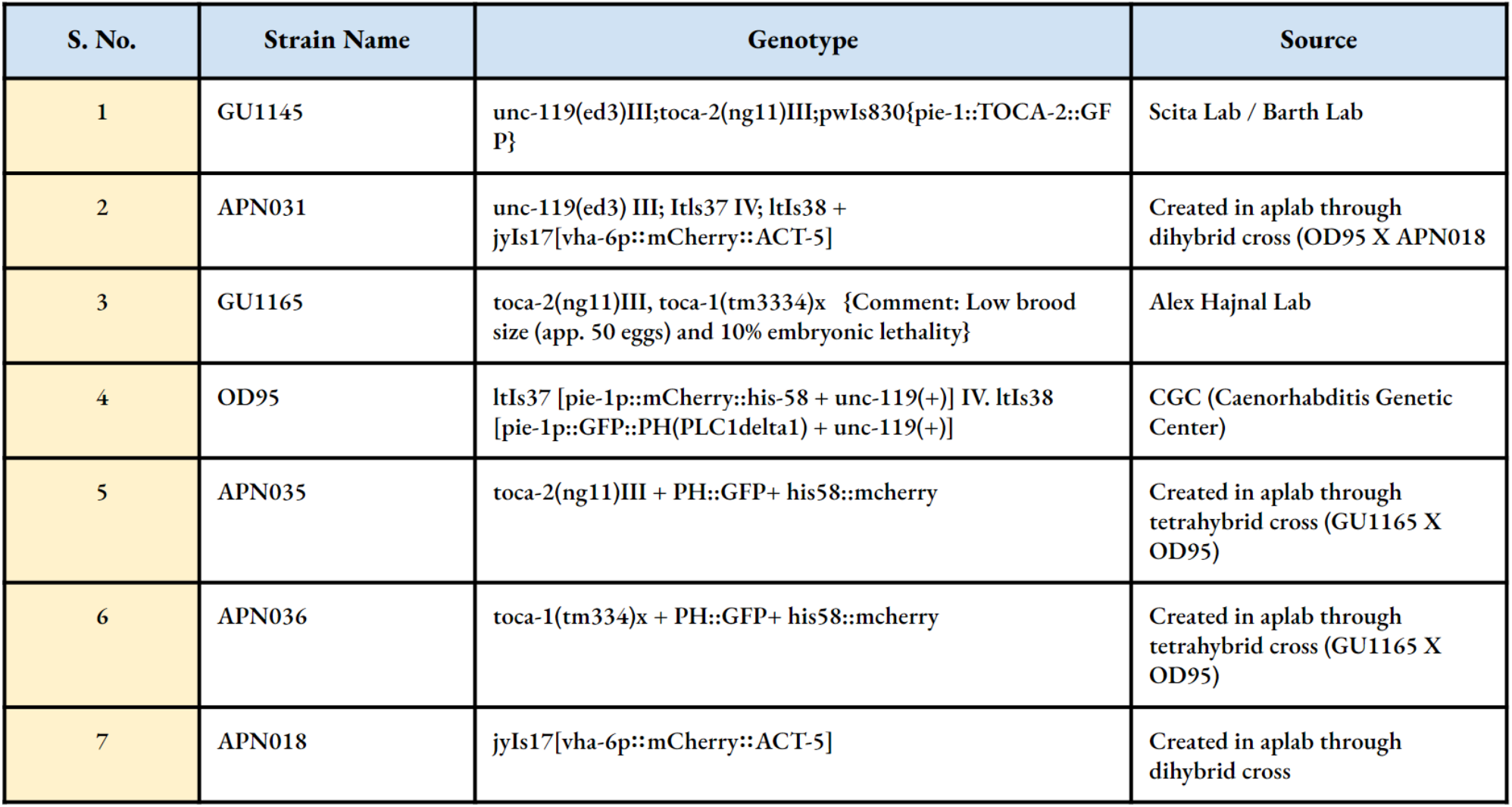

**Table S2.**
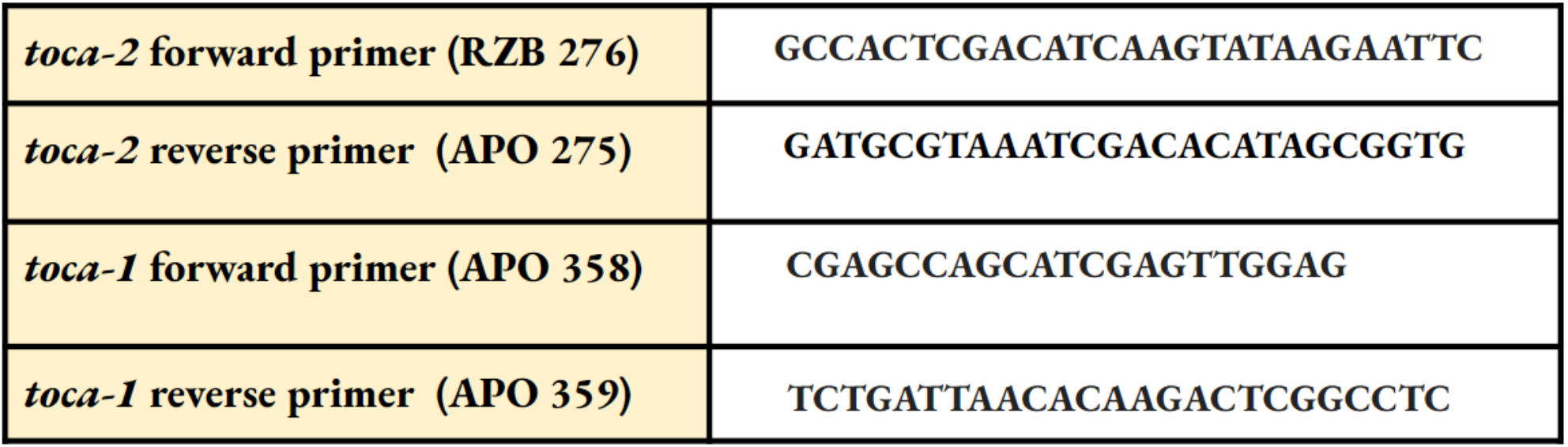

**Figure S1:**
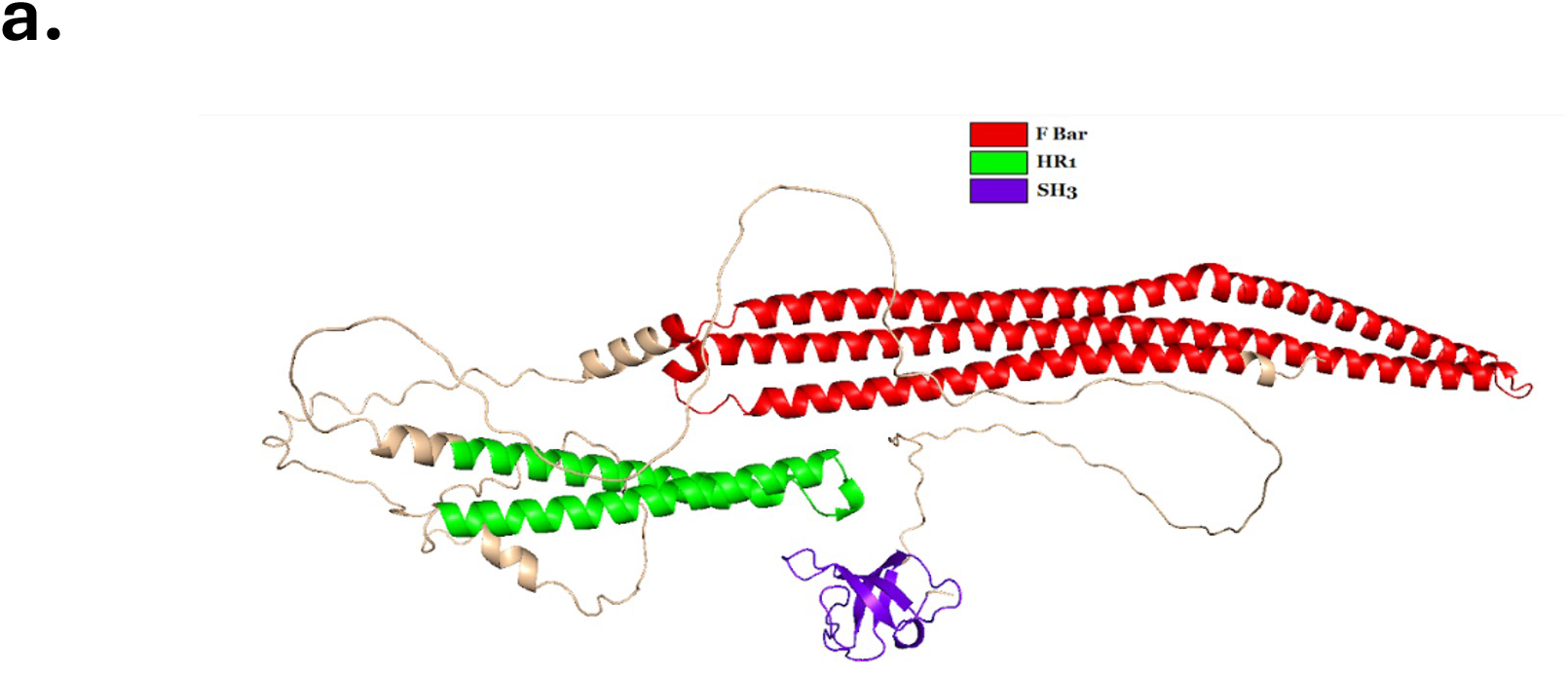
TOCA-2 contains a curvature sensing F-BAR domain at the N-terminus, an HR-1 domain that binds to CDC-42 and an N-WASP binding SH3 domain at the C-terminus.

**Figure S2.**
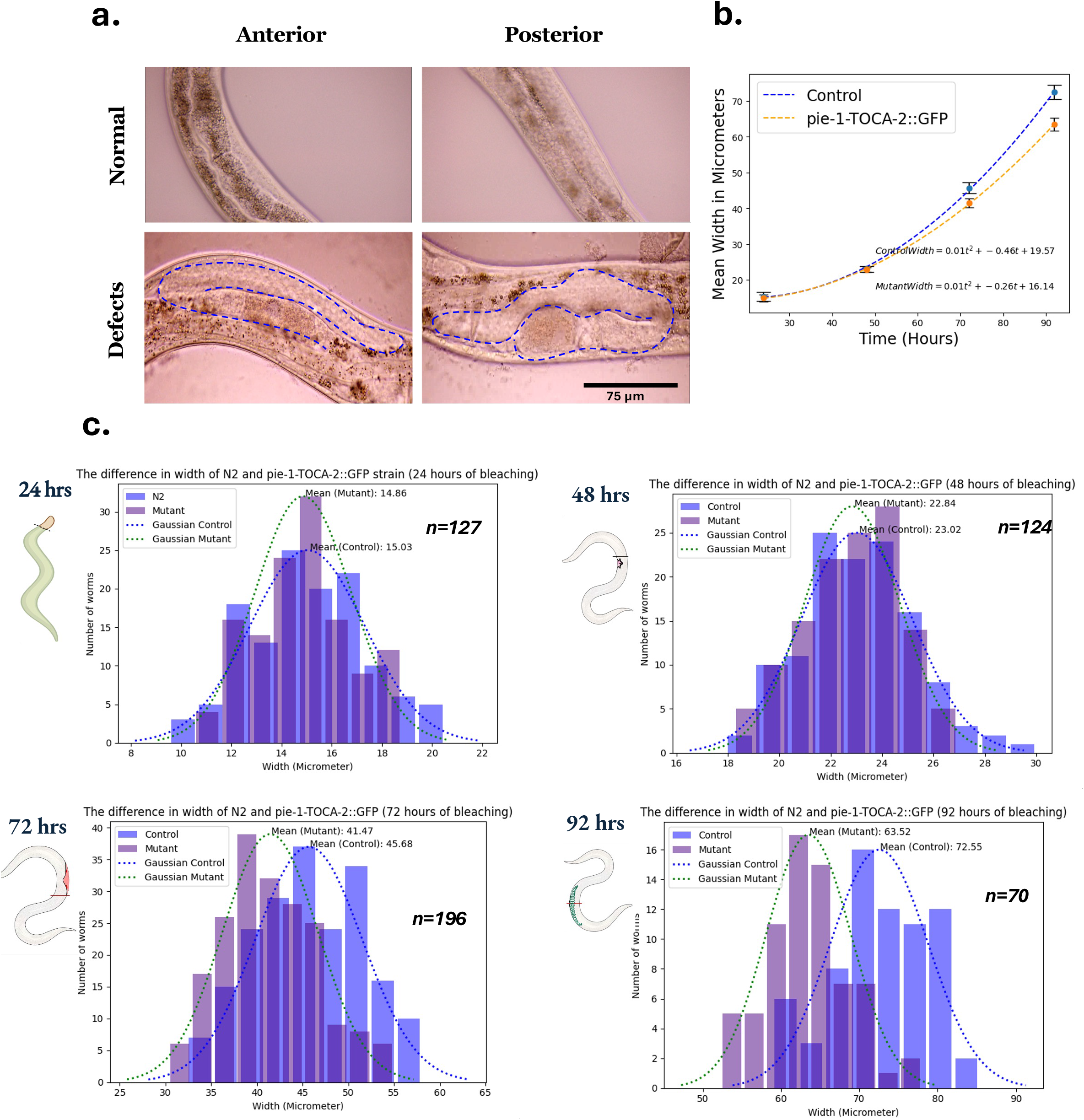
**a**. The initial images of defects observed in the mutant showing germ line-specific expression of TOCA-2. **b**. Cumulative graph showing time-course width analysis between control and *pie-1::*TOCA-2::GFP. c. Plot showing the histogram of the width of *pie-1-*TOCA-2::GFP and control at each time point.

**Figure S3.**
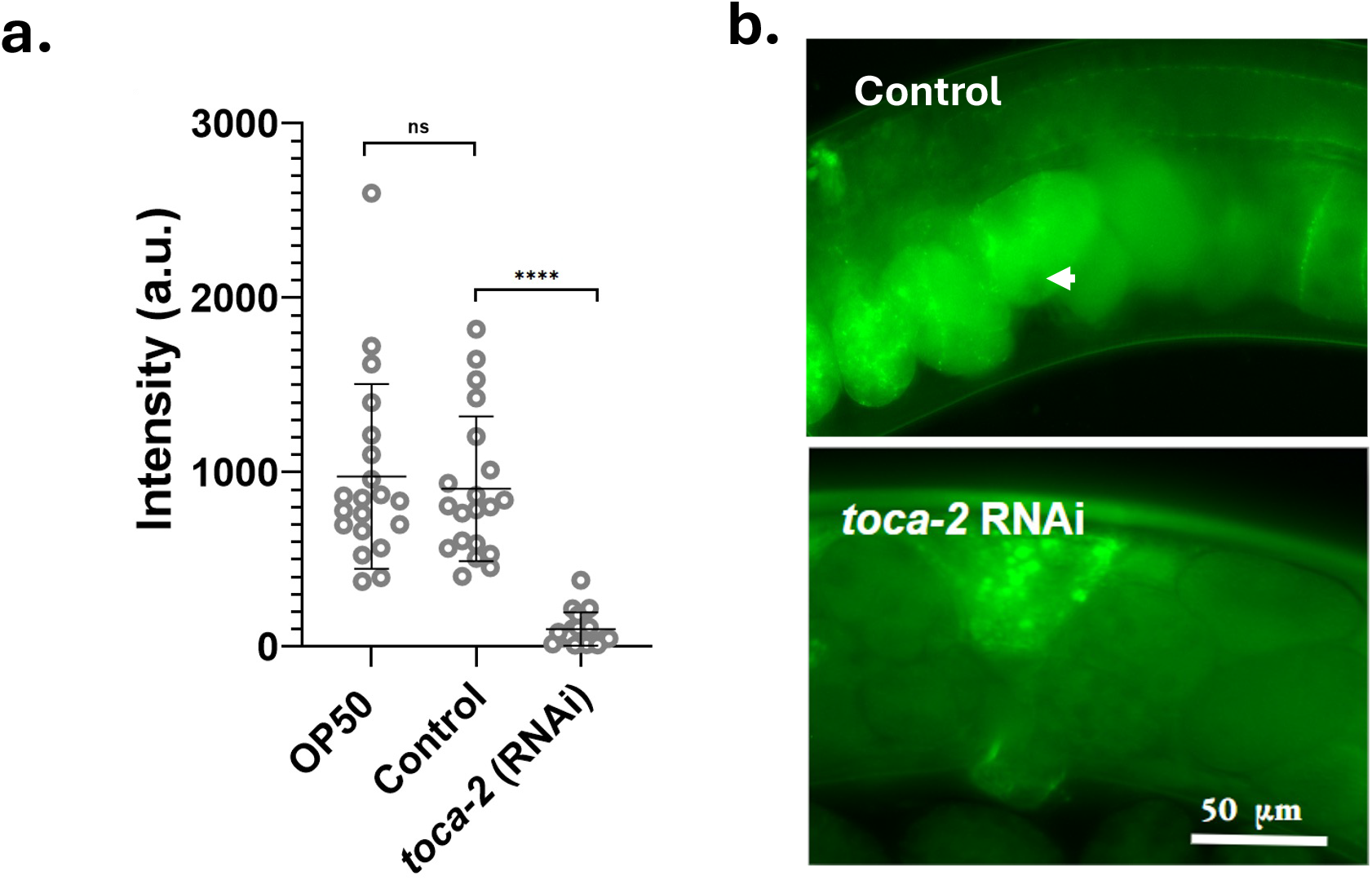
a. Intensity analysis after toca-2 *RNAi* for two consecutive days b. Micrographs representing intensity of TOCA-2 ::GFP after RNAi-mediated depletion of TOCA-2

**Figure S4.**
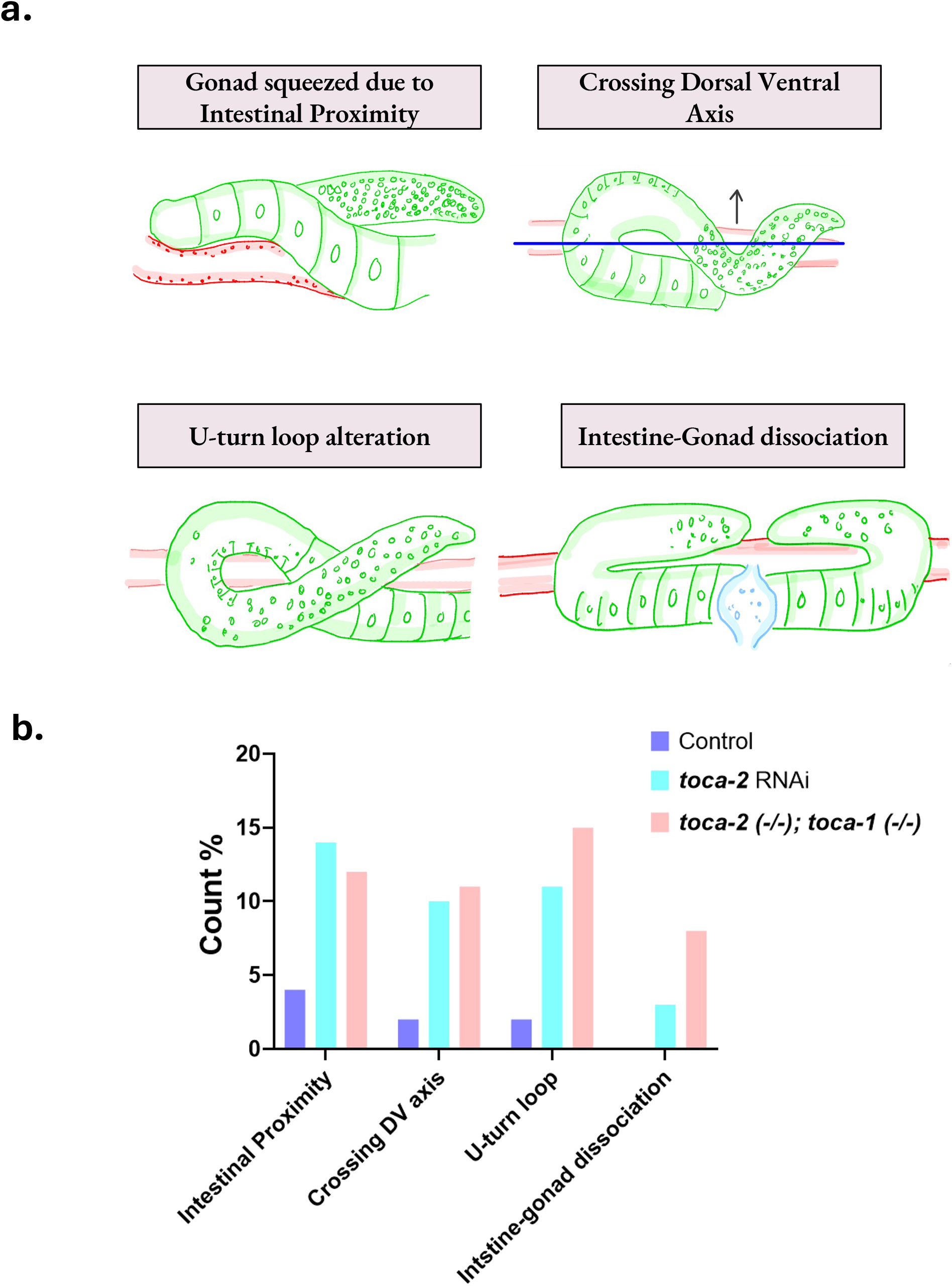
Characterization of the defects observed in the control, *toca-2 (RNAi)*, and *toca-2 (-/-) :: toca-1 (-/-)*. ‘Uturn loop morphology’ is the most prominent defect in TOCA-2 mutants.

**Figure S5.**
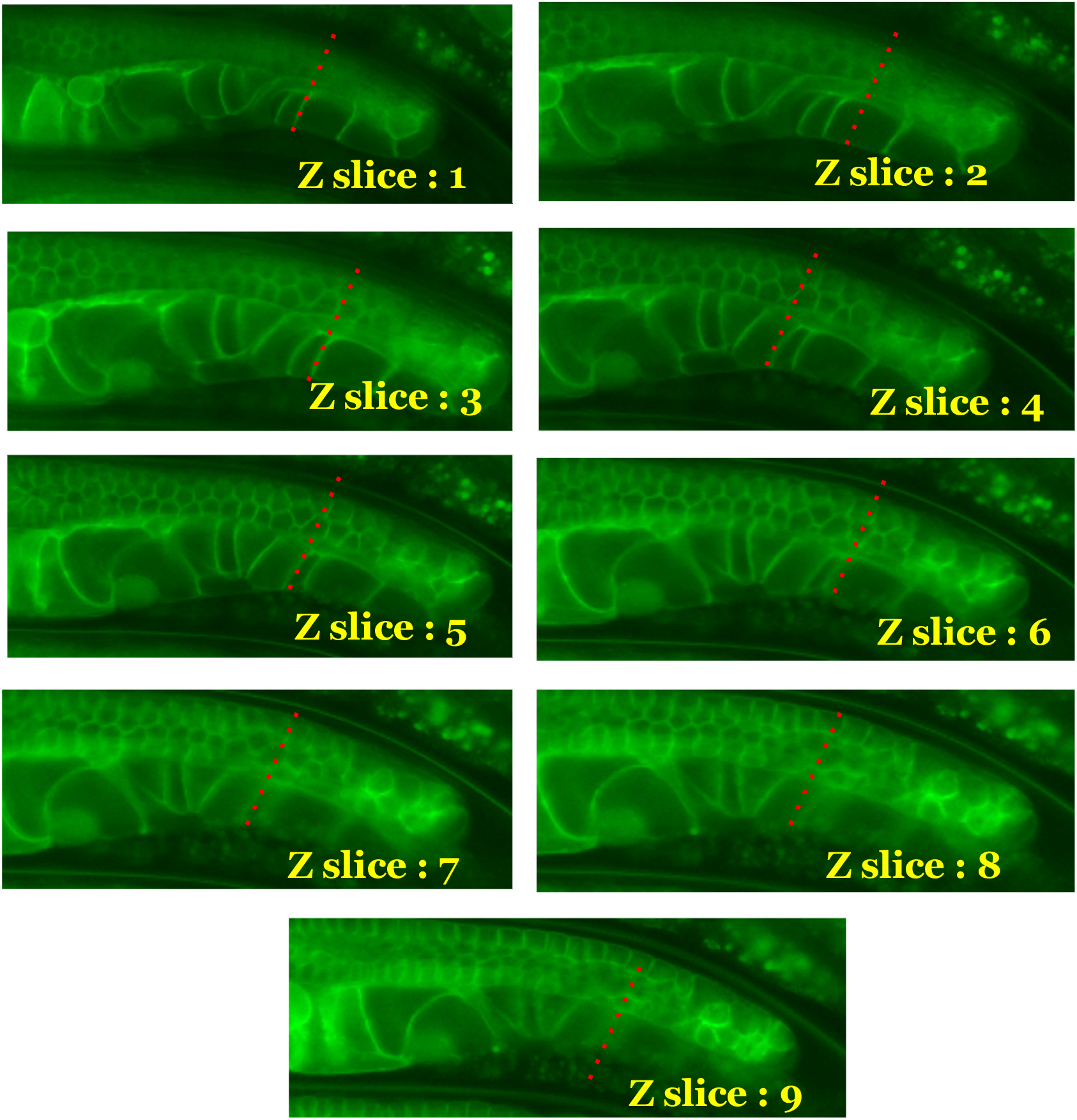
Analysis to assess the discrepancy in gonad width due to variations in the z-plane

**Figure S6.**
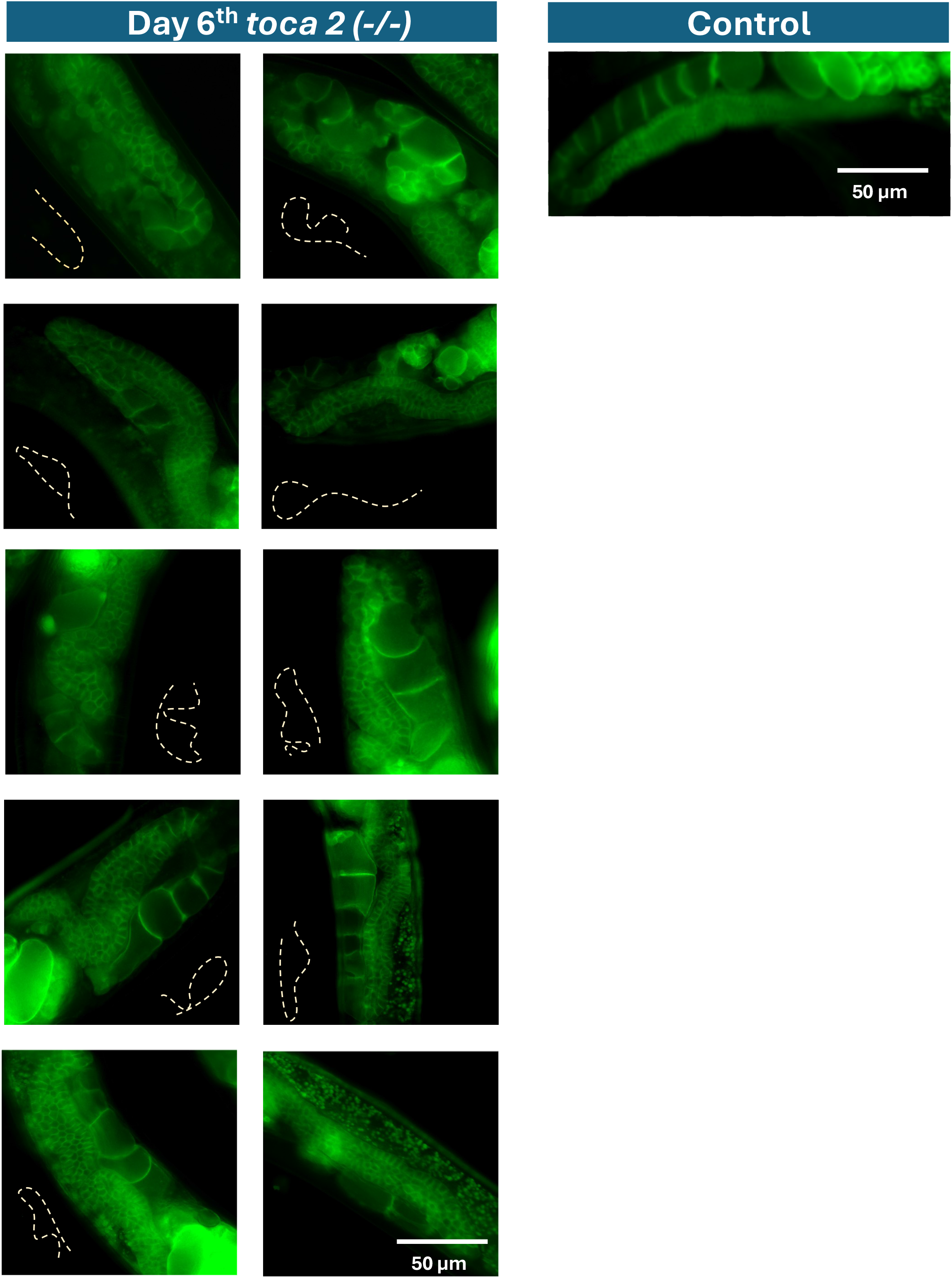
In the *toca-2(null)* mutant, phenotype of gonad atrophy was observed, which was characterized by a significant decrease in gonad length, with the most pronounced shrinkage noted towards the dorsal side.

**Figure S7.**
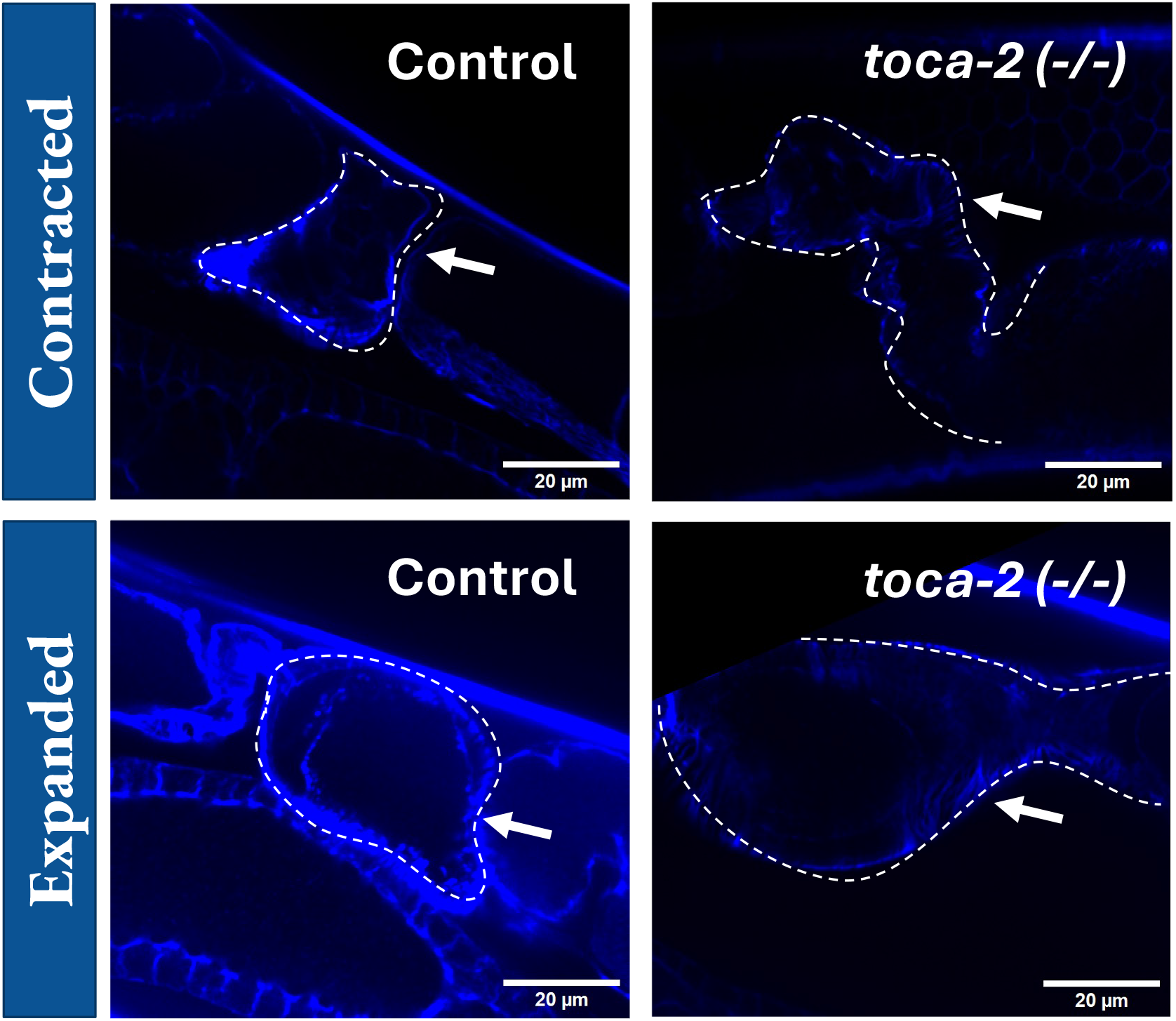
The structure of the spermatheca in the *toca-2(null)* mutant exhibited a deviation from the control. The first panel depicts the spermatheca’s structure in a contracted state, while the second panel displays the image of the expanded spermatheca (with the presence of oocytes).

**Figure S8.**
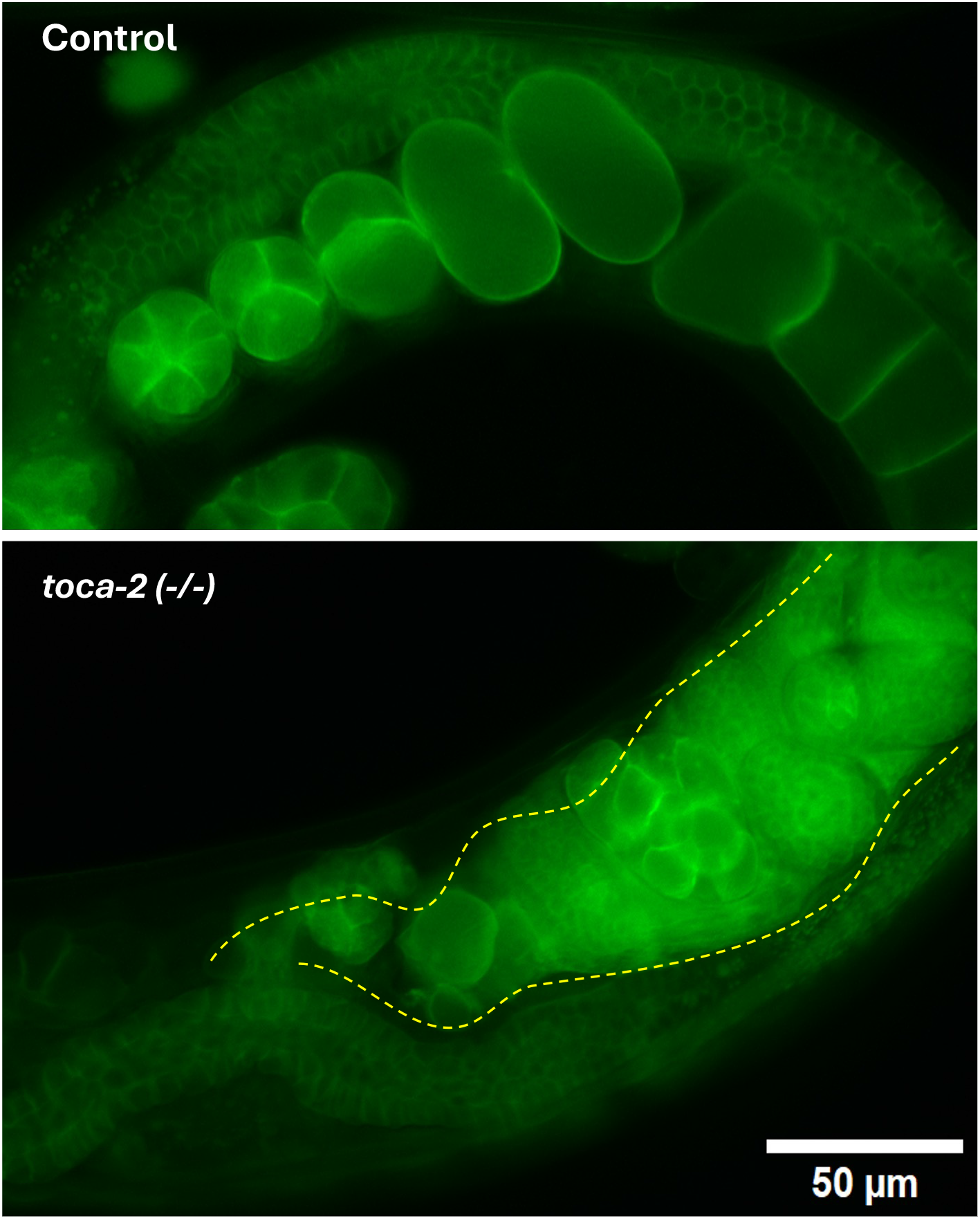
Teratoma-like structures observed in the gonads of *toca-2(null)* animals

